# Hornbills Adjust Response Speed According to Solvability of Patterned-Strings Problems

**DOI:** 10.1101/2025.06.25.661464

**Authors:** Chunyun Xue, Elias Garcia-Pelegrin

## Abstract

Cognitive adaptations for processing causal information are fundamental to flexible problem-solving across species. Patterned-string tasks offer a classic means–end problem, requiring subjects to identify the functional connection between an action and its distal outcome. This study investigated patterned-string pulling in East Asian hornbills, a cognitively underexplored avian taxon, to assess their problem-solving abilities and underlying cognitive mechanisms. Nine hornbills were presented with three string configurations of increasing complexity: parallel (baseline), broken (contact/no-contact), and crossed strings. Subjects reliably selected the correct string in the baseline and contact conditions, demonstrating sensitivity to physical continuity cues. In contrast, accuracy declined significantly in the crossed condition, which required recognition of spatial displacement between the baited and accessible string ends. Reaction time (RT) analyses revealed prolonged latencies in the crossed condition, likely reflecting increased cognitive demands or limited habituation. Individuals who failed often adopted suboptimal strategies, such as proximity or side biases. We also observed lateralisation effects, with a left-side RT advantage, suggesting hemispheric specialisation. These findings support the hypothesis that task complexity modulates both decision accuracy and response speed, highlight RT as a useful proxy for interim cognitive processing, and establish hornbills as a valuable comparative model for studying causal cognition in birds.

## Introduction

Understanding the causal relationship between actions and their outcomes is crucial for effective problem-solving. This knowledge is typically defined as the cognitive ability to comprehend a series of actions (the “means”) required to achieve the end goal (Huber & Gajdon, 2006). Piaget (1952) suggested that solving a means-end problem is one of the first significant landmarks of ’*intelligent behaviour* ’, which is believed to emerge at approximately the age of 5-8 months and continue to refine in early human development (e.g., Casler & Kelemen, 2005; Lobo & Galloway, 2008; Munakata et al., 1997; Nielsen, 2006; Rat-Fischer et al., 2012; Willatts, 1999; Wimpenny et al., 2009). The means-end understanding is often tested through the patterned string-pulling task, where one or more strings are arranged in patterns such as: *parallel*, *perpendicular*, or more complex arrangements (e.g., *slanted* in parallel, *crossed* together), with a reward attached to one of the strings at an out-of-reach distance (see Jacobs & Osvath, 2015, for a review). The subject must retrieve the reward by first recognising the exact string with the reward attached and then pulling until it is within reach. While developmental inquiries employing patterned-string tasks frequently investigate the longitudinal development of means-end understanding and goal-directedness in humans (Frye, 1991); studies in non-human animals endeavour to map phylogenetic differences in this problem-solving ability across species (for reviews, see: Jacobs, 2018; Jacobs & Osvath, 2015). To date, the patterned string-pulling tasks have been administered to over 160 species, and most prevalently tested in avian species under the orders of songbirds (*Passerines*; e.g., Baciadonna et al., 2022; Bagotskaya et al., 2012; Heinrich, 1995; Obozova et al., 2014; Taylor, Medina, et al., 2010; Taylor et al., 2012) and parrots (*Psittaciformes*; e.g., Bastos et al., 2021; Gaycken et al., 2019; Krasheninnikova, 2013, 2018; Krasheninnikova et al., 2013; Pepperberg, 2004; Schuck-Paim et al., 2009; Wakonig et al., 2021).

A growing body of evidence suggests that corvids and parrots possess cognitive abilities in problem-solving that are comparable to those of large-brained mammals such as great apes (Emery & Clayton, 2004). This has been evidenced in both field and laboratory studies, where corvids and parrots have shown remarkable flexibility in various cognitive tasks, including social vocal learning (e.g., Bradbury & Balsby, 2016; Pepperberg & McLaughlin, 1996; Pepperberg et al., 1997), inhibitory control (e.g., Koepke et al., 2015; Miller et al., 2019, 2023), mental time travel (e.g., Clayton & Dickinson, 1998; Clayton et al., 2003), and tool-use (e.g., Bird & Emery, 2009; Clayton & Dickinson, 1998; Emery, 2006; Taylor, Elliffe, et al., 2010; Weir & Kacelnik, 2006; Weir et al., 2002). The higher-order cognition is suggested to be a product of their densely packed neurons in the brain, with some corvids and large-sized parrots being found to exhibit equivalent or even greater neuronal densities in the forebrain area compared to monkeys (Olkowicz et al., 2016; Sol et al., 2022). Particularly, within the forebrain area lies a segment known as the *nidopallium caudolaterale*(NCL), believed to share functional homology with the mammalian prefrontal cortex and support executive functions in birds (Divac et al., 1985; Güntürkün, 2005; Güntürkün & Bugnyar, 2016).

However, it is worth noting that, besides species under the *Passerines* and *Psittaciformes* orders, the cognitive abilities of other large-brained birds are still under-represented in the current empirical literature. For instance, species of the hornbill family (*Bucerotidae*, order *Bucerotiformes*) are found to have enlarged brain size, with their relative brain size surpassing that of some parrots (Iwaniuk et al., 2005). Hornbills mostly inhabit tropical and subtropical regions in Africa and southern Asia, living an average life expectancy of 20 years, while larger individuals may enjoy an increased life expectancy of 50 years (Kemp, 2007). Most hornbills have an omnivorous diet, consisting of seeds, insects, small animals, and a high prevalence of fruits. In recent years, East Asian hornbills, notably the oriental pied hornbills (*Anthracoceros albirostris*), originally confined to nesting in tree cavities in tropical regions, have displayed high adaptability to urbanization by breeding and nesting in semi-urban areas with artificial infrastructure (e.g., Datta & Rawat, 2003; James & Kannan, 2007; Kalina, 1989; Kozlowski et al., 2015; Loong et al., 2021; Shukla et al., 2015). This adaptive success is often thought to be supported by advanced cognitive capacity (Garcia-Pelegrin et al., 2022; Sol et al., 2002). More recent empirical findings reveal that oriental pied hornbills, *Anthracoceros albirostris*, can exhibit full Piagetian object permanence and complex task specific problem-solving (Garcia-Pelegrin, 2025; Yao & Garcia-Pelegrin, 2024), as observed in parrots and corvids (e.g., Pepperberg et al., 1997; Zucca et al., 2007), further confirming the enhanced cognitive skills of this species. Hence, it is of great interest to explore whether hornbills can demonstrate similar cognitive phenomena in problem-solving contexts, such as the means-end understanding.

Many cognitive processes contribute to solving means-end patterned-string problems, including associative/reinforcement learning (RL), causal reasoning, and insight (Emery & Clayton, 2004; Heinrich, 1995; Heinrich & Bugnyar, 2005; Huber & Gajdon, 2006; Jacobs, 2018; Jacobs & Osvath, 2015). These theoretical frameworks differ in the complexity of information processing they entail. The *associativist* perspective posits that problem-solving emerges through trial-and-error learning and the formation of associations between actions and rewards, without requiring an understanding of the underlying causal mechanisms. As such, more errors during the task are expected if there are “unlearned” changes or increased complexity in the task context (e.g., Taylor et al., 2012). In contrast, the causal reasoning account suggests a more flexible cognitive mechanism by which an animal, either spontaneously or through experience, infers the causal relationship between actions and outcomes, potentially enhancing behavioural adaptability in dynamic environments (Huber & Gajdon, 2006; Krasheninnikova, 2018). The insight hypothesis, often linked to imagination, proposes that an animal not only understands causality based on prior experience but also, upon reaching a threshold of accumulated knowledge, can mentally simulate future actions to generate a solution (Emery & Clayton, 2004; Epstein et al., 1984). However, empirical literature sometimes presents these theories as parallel rather than rival explanations, a distinction that lies beyond the scope of the present discussion on means-end problems in avian string-pulling (e.g., Colin & Belpaeme, 2019; Wasserman & Castro, 2022).

Current evidence from patterned-string experiments does not support models proposing complex cognitive mechanisms. For example, experiments using a vertical patterned-string apparatus showed that corvids and parrots, such as New Caledonian crows (Taylor, Elliffe, et al., 2010; Taylor et al., 2012), Goffin’s cockatoo (Wakonig et al., 2021), keas (Werdenich & Huber, 2006), and galahs (Krasheninnikova, 2013) succeeded in a two-string discrimination task, but chose at random when the task content was altered, such as in a crossed-string task (i.e., two strings are crossed so that the end of the baited string is farther away than the end of the unbaited string). The failures in a crossed-string task may indicate a lack of understanding of the causal relationship between the strings and rewards. Moreover, Taylor, Elliffe, et al. (2010) showed that when visual feedback was restricted, crows interacted with the strings with significantly more errors and less efficiency, indicating an absence of ’insight’ but solely reliance on perceptual feedback during testing. Therefore subjects may rely on simple strategies, like associative learning between visual feedback of string proximity and the reward, rather than comprehending the underlying physical relationships or attempting to ’plan’ for future actions (but see, Garcia-Pelegrin, 2025). However, research employing a horizontal setup observed even though some subjects demonstrated little knowledge of the physical relationships of strings in the *crossed* version, they showed relative ease in discriminating between an intact and another string broken in the middle with a visible gap (i.e., *contact/no contact* version), where both strings are baited (e.g., Johnsson et al., 2023; Krasheninnikova et al., 2013; Obozova et al., 2014; Wang et al., 2019).

From the avian string-pulling empirical literature, three potential explanations emerge for the mixed results in patterned-string-pulling studies: (i) *partial causal understanding*, wherein subjects may possess only limited comprehension of physical concept; (ii) *proximity rule*, where subjects rely solely on linear visual feedback (i.e., proximity) of the rewards; and (iii) *visual affordances*, states that the subjects may be more sensitivity the physical connection of a string by default or due to training (e.g., Jacobs, 2018; Jacobs & Osvath, 2015; Obozova et al., 2014). The *partial understanding* account and the *proximity rule* are often linked together to explain the frequent failures observed under the *crossed*-strings paradigm. The underlying logic is that subjects’ “*partial* knowledge” of the spatial affordances of the strings, combined with a strong bias toward the more proximal end attached to a reward, makes it difficult for them to disentangle the *crossed* pattern. However, we identify potential ambiguity in these accounts. First, the concept of “*partial* understanding” operates primarily at a meta-cognitive level and fails to address the specific cognitive barriers preventing full causal reasoning about the strings. Additionally, measuring “partial understanding” is methodologically challenging, rendering it potentially unfalsifiable. Similarly, the *proximity rule* explanation may lack generalisability, as it seems not to account for why some birds perform randomly under *crossed*-string conditions (Hofmann et al., 2016; Krasheninnikova et al., 2013) but do not choose randomly in *contact/no-contact* tasks, despite proximity being their primary strategy (Wang et al., 2019). In contrast, the *visual attention* hypothesis appears better supported by empirical evidence. For instance, some parrots succeed only when *crossed* strings are marked with visually distinctive colours but fail when these cues are removed (Krasheninnikova et al., 2013).

Another limitation of current empirical approaches is the reliance on response accuracy as a single metric, which may be insufficient to fully capture the perceptual and cognitive processes underlying decision-making in patterned string tasks. In addition to such discrete outcome measures, temporal variables, particularly response latency (RT)—have traditionally been interpreted in behavioural and cognitive research as proxies for mental computation. These latencies can be shaped by factors such as task difficulty, response accuracy and heuristics (e.g., Diedrichsen et al., 2010; Ratcliff, 1978, 2002; Wong et al., 2017). Similar assumptions are found in avian cognition and neuroscience, where RT is often taken as an index of attentional and decisional processes (e.g., David et al., 2014; Nebel et al., 2019; Sridharan et al., 2014). For instance, Sridharan et al. (2014) showed that domestic chickens performed more quickly and accurately in a cued spatial discrimination task, indicating enhanced attention and stimulus prioritisation. Similarly, Nebel et al. (2019) found that feral pigeons responded more slowly to predator silhouettes under dim or low-contrast visual conditions, suggesting increased perceptual demands. However, recent work in human motor learning and decision-making has challenged the notion that RT purely reflects cognitive effort, showing instead that it can also be shaped by habitual or experience-based biases independent of deliberative processing (e.g., Wong et al., 2016, 2017).

Building on this theoretical background, the present study investigates whether hornbills’ response latency varies across different patterned-string tasks. We hypothesise that if their choices are guided by simple proximity-based heuristics that require minimal cognitive effort, reaction times (RTs) should remain relatively consistent across tasks. In contrast, if choices involve active visual assessment of string configurations, RTs are expected to vary depending on task-specific features such as spatial complexity. To assess the influence of experience on performance, we adopted a learning-based framework: subjects were first trained on a simple condition (e.g., two-string discrimination), then exposed to a more complex and unfamiliar task (e.g., crossed strings), and finally re-tested on the original simple task. If hornbill problem-solving is influenced by prior experience, we expect to observe carry-over effects in both accuracy and latency, reflecting generalisation or interference from the preceding task. Alternatively, if performance remains stable before and after exposure to the novel condition, this would suggest that decision-making is governed by task-specific strategies rather than broader learning-based modulation

## Methods

### Subjects and Housing

A total of nine hornbills participated in the current study. Eight of them belong to the genus *Rhyticeros*, including six oriental-pied hornbills (*Anthracoceros albirostris*; three females named Chiku, Oris, and Olivia; 3 males named Chika, Sam, and Oscar), one plain-pouched hornbill (*Rhyticeros subruficollis*, female named Harriet), and one Papuan hornbill (*Rhyticeros plicatus*, female named Pepper), all of which are native to Southeast Asia. Additionally, one trumpeter hornbill from the genus *Bycanistes* (*Bycanistes bucinator*, female named Sadie), native to Africa, was also tested.

All subjects are part of the Mandai Wildlife Reserve collection and are housed either independently (if unpaired) or in pairs in aviaries measuring approximately 6m in width, 4m in length, and 3m in height. The hornbills were either rescued from illegal wildlife trafficking or had suffered severe injuries before fledging in the wild; hence, their precise ages are unknown, though their estimated age range is 5 to 10 years. None of the hornbills were food-deprived during training or testing. They were fed a maintenance diet of fresh fruits and nutrient pellets and had *ad libitum* access to water.

### Procedures

The experimental protocol was reviewed and approved by the National University of Singapore’s Institutional Animal Care and Use Committee (IACUC protocol number R23-0737) and the Mandai Wildlife Reserve research panel.

### Experimental Design

We tested three different patterned-string conditions in the current experiment: parallel (*baseline*), *crossed*, and *contact* (see Figure 1a for a simple visual illustration of the string patterns). None of the subjects have been exposed to patterned-string tasks in prior to training. The experiment adopted a block design, with a total of 9 blocks grouped into 3 experimental sessions. Each session was designed to follow a predetermined sequence: *pre-test baseline block* → *test block* → *post-test baseline block* to monitor changes in choice accuracy and RT. In each session, parallel-patterned strings were presented to the subjects in the *pre-test baseline block* and *post-test baseline block*, while in the *test block*, we manipulated the string patterns to be *crossed*, *contact*, and *baseline* control. Each block contained 20 consecutive trials. In each trial, the experimenter first centered the subjects in a middle position using the food reward, then the experimenter manually baited the distant end of the strings (as shown in Figure1b), with the side of the baited string pseudo-randomised. Afterwards, the experimenter pushed the reachable ends into the subjects’ enclosure for them to make decisions. Throughout this process, we maintained maximal symmetry in hand movements to avoid any potential misleading effects from gestural signaling.

**Figure 1.**
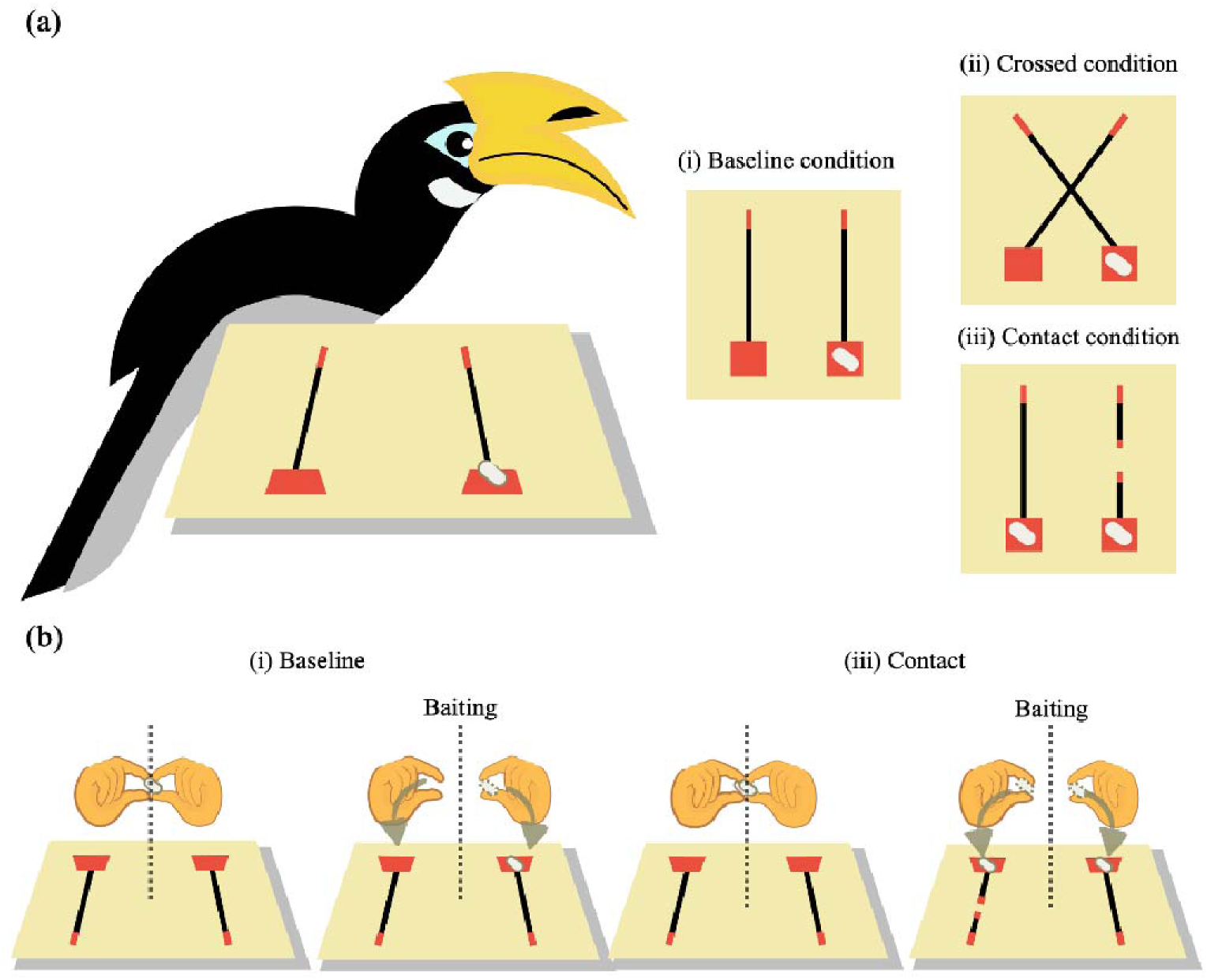
A simple visual illustration of the patterned-string layout and baiting process. *Note:* The string patterns of the control and experimental conditions are listed in panel (a). In (i) *baseline*, two strings are placed parallel to each other and orthogonal to the side of the board adjacent to the subject. Similarly, in (ii) *crossed*, the strings are crossed in the middle. In (iii) *contact*, one of the strings is broken with a visible gap (approximately 5 cm in length) in the middle. Panel (b) illustrates the hand movement during baiting. In (i) *baseline*, the experimenter first centred the subject with a food reward and then baited one of the strings. In (iii) *contact*, the experimenter baited both strings. Hand movements were kept maximally symmetrical to avoid gestural following. The sides of the baited strings (or the broken strings) were pseudo-randomised across the trials within one block.

### Test Apparatus

During the experiment, the subject faced directly to a 35cm *×* 20cm wooden board placed adjacent to it’s enclosure. We used the string in 30cm length, with one square 2cm *×* 2cm container tied to one end in order to carry food reward (see Figure1a). The ends of two strings were placed in approximately 15cm apart from each other. For the *contact test block* specifically, we left a 5cm gap between the broken strings. The food rewards we used were either avian food pellets or fresh fruits sliced into 5mm-10mm cubes derived from subjects’ daily diet portions

### Pre-training

Subjects were initially allowed to familiarise themselves with the single-string setup, where one string with a reward attached to the distant end was placed in front of them. Testing sessions started only after subjects successfully pulled the single string and obtained the reward in 10 consecutive trials. All subjects passed the single-string phase within 15 trials.

### Testing

Subjects proceeded to the testing phase once they passed the single-string training. The order of sessions tested for each subject is listed in Supplementary material A, Table S1. Most subjects received one session per test day. If any subjects were tested with more than one session per test day, the inter-session interval was set for at least 4 hours. The average duration of one experimental session was 402.51 seconds (*SD*: 76.41 seconds). The variation in session duration was due to the subjects’ occasional loss of interest in the task. If a subject disengaged from the task for over 5 minutes, the experimenter discontinued the session and resumed after 10-15 minutes or on the next test day.

Each session started with a pre-test baseline block to habituate the subjects to the task. If a subject scored lower than 14 out of 20 in the habituation block, that testing session was aborted and would resume after 10-15 minutes or on the next test day. Otherwise, the subjects proceeded to the test block, followed by the post-test baseline block. The maximum inter-block duration was around 120 seconds. Although the aim was to conduct 20 trials per block, some subjects had 1-3 additional trials for attention or side-preference corrections. The average number of trials per block was 20.33 (*SD*: 0.71).

### Data

All the testing sessions were video-recorded using a GoPro action camera (model: HERO 11) and coded using BORIS version 8.22.16 (Friard & Gamba, 2016). In every experimental trial, the experimenter’s baiting time (in ms), the side of the baited string (L: left *vs* R: right), the side of the chosen string (L: left *vs* R: right), subjects’ choice outcome (correct *vs* incorrect), and subjects’ reaction time (RT; in ms) were coded (for the data coding, see Supplementary material B). We further processed the RT data for analyses by filtering out the extreme anomalies using Tukey’s methods (Tukey, 1977, for details, see Supplementary material C).

### Analysis

All analyses were conducted using R statistical software version 4.3.1 (R Core Team, 2023). Exact binomial tests with an expectation of 0.5 were performed on choice and chosen side individually to examine the performance and biases of string-pulling in each test block. We further fitted generalized linear mixed models (GLMMs) on the choice accuracy and RT data using the “glmm” package (Knudson, 2022).

We conducted GLMMs to assess the fixed effects of session (3 levels: crossed *vs* contact *vs* baseline), condition/block (3 levels: pre-test baseline *vs* test *vs* post-test baseline), side of the baited string (2 levels: left *vs* right), side of the chosen string (2 levels: left *vs* right), baiting duration (measured in *ms*), trial sequence per test block, and the interaction of session by condition. “DHARMa” package was employed to check if model assumptions were met (Hartig, 2022). In order to meet the model assumptions for RT analysis, we transformed the RT by taking the base-10 logarithm, as empirically observed RTs usually follow log-normal distributions (Salinas Ruíz et al., 2023; Ulrich & Miller, 1993).

For the binomial GLMM on choice accuracy, we added by-subject random intercepts, while for the GLMM on RT, we further added by-subject random slopes of *session* and *condition*. Note that we removed the fixed effect of *baited side* in both GLMMs to ensure model conversion. The ANOVA tables with type III method were derived from the GLMMs using the “car” package (Fox & Weisberg, 2019) since some of our predictors are multi-level, and *p*-values, *F* -statistics, and χ^2^ statistics are reported in the **Results** section. Post-hoc pairwise comparisons were conducted using the “emmeans” package (Lenth, 2024) with *Bonferroni-Holm* methods to adjust *p*-values.

## Results

### Accuracy

As shown in Figure 2, subjects performed consistently across the first two blocks, selecting the correct string significantly above chance in the *baseline* session. In the *contact* session, 5 out of 9 hornbills (Chiku, Harriet, Olivia, Oris, and Sam) also chose the correct string significantly above chance during the *test* block. In contrast, no subjects showeda significant preference for the correct string in the *crossed* session; instead, 4 subjects (Chika, Oris, Oscar, and Pepper) showed a significant preference for the incorrect string.

**Figure 2.**
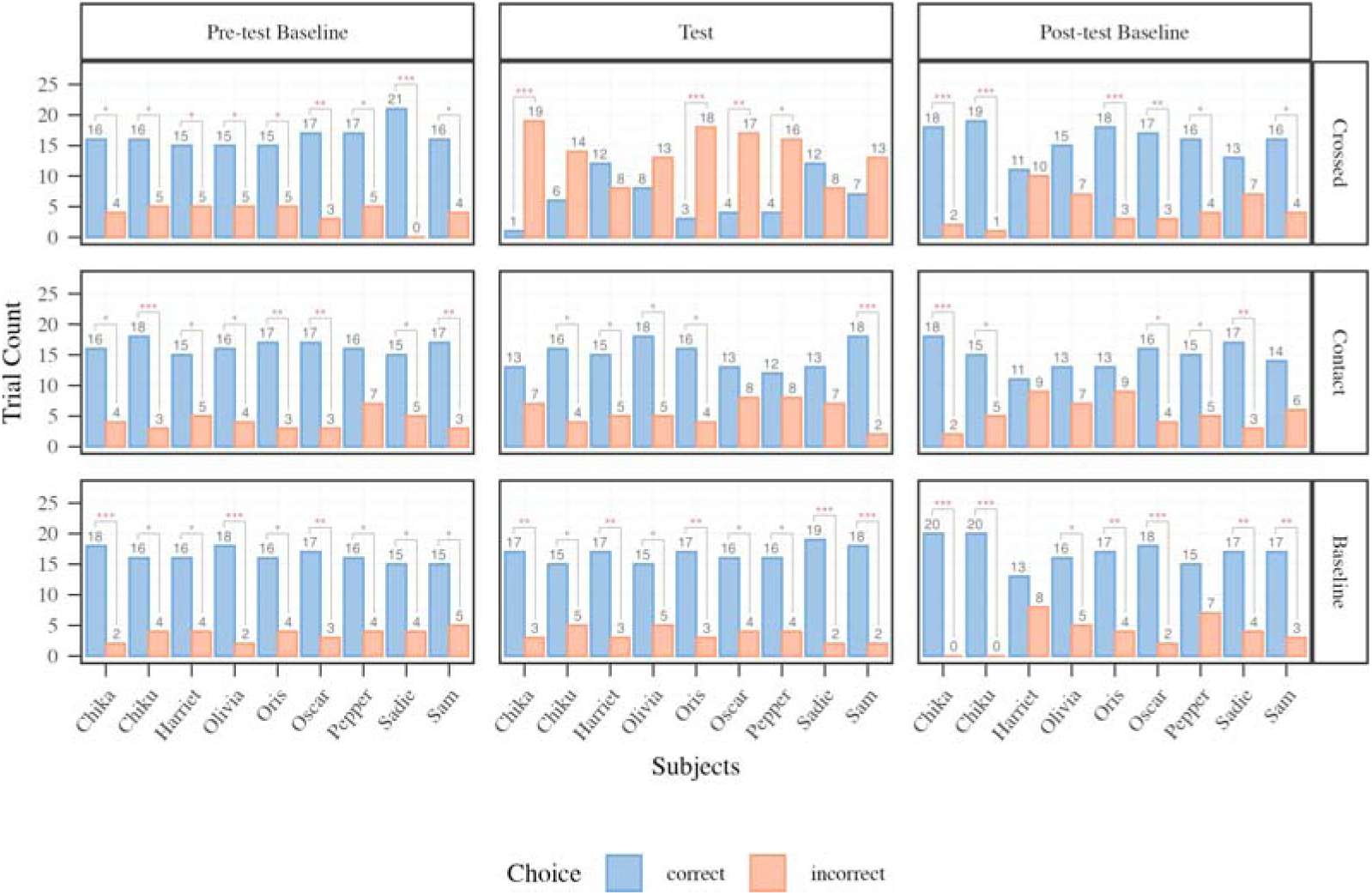
A barplot summarises individual performance per block. *Note:* the number of correct/incorrect trials per block is summarised above each coloured bar. Binomial significance levels are denoted by annotations coloured in red: \**p* < .05. \*\**p* < .01. \*\*\**p* < .001.

Further binomial exact tests on side preferences revealed that 4 subjects (Chiku, Olivia, Sadie, and Sam) developed a significant side bias during the test block of the *crossed* session. Of these, Olivia and Sadie maintained this bias in the subsequent baseline block. In the *contact* session, only Sadie showed a significant side bias during both the pretest baseline and test blocks, while Olivia and Oris developed a bias only in the post-test baseline block. No subjects exhibited a significant side preference during the *baseline* sessions. For details on binomial significance levels, see Figure 3.

**Figure 3.**
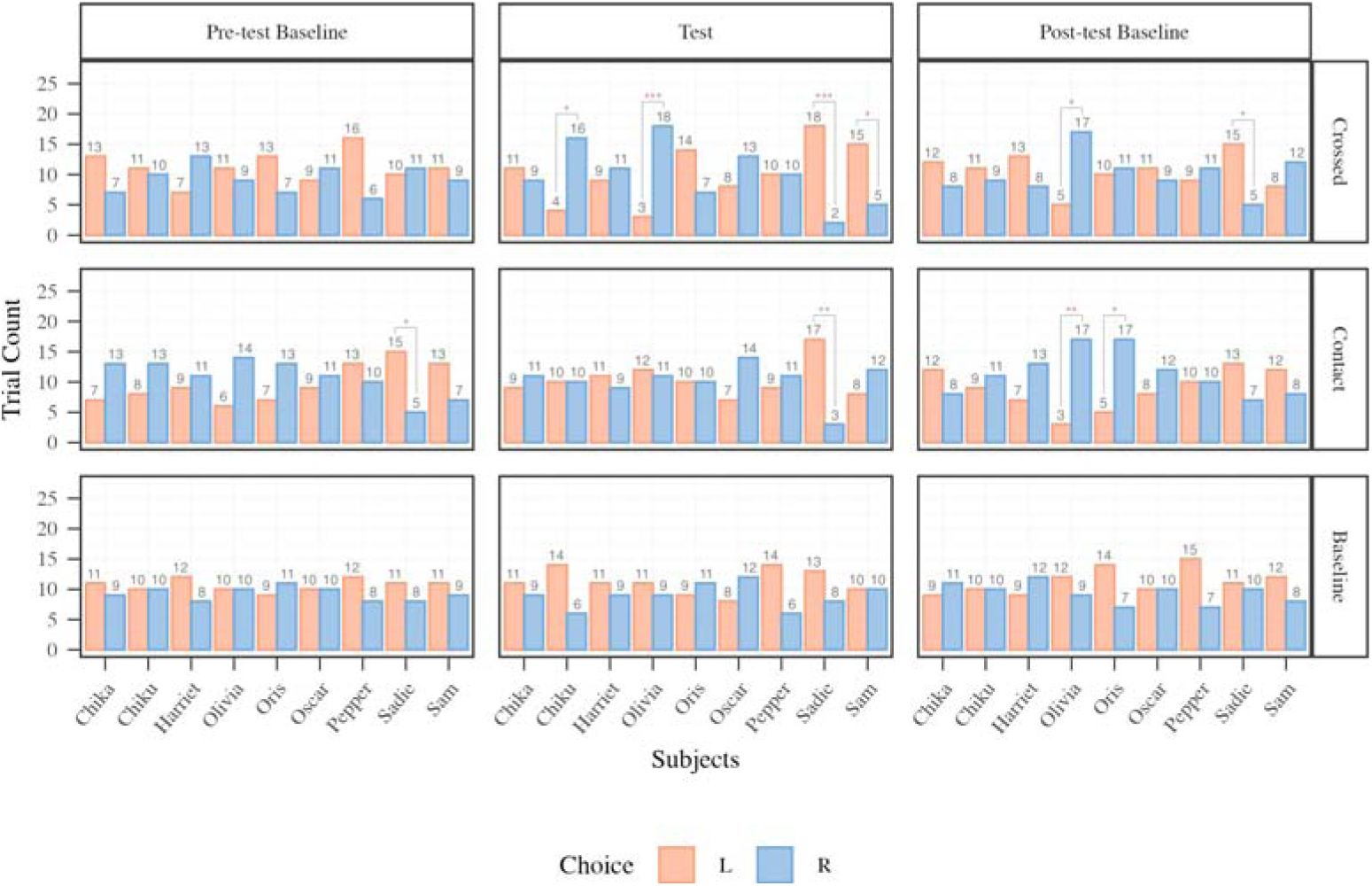
A barplot summarises individual side preference per block. *Note:* the number of sides (right vs left) of the chosen string per block is summarised above each coloured bar. Binomial significance levels are denoted by annotations coloured in red: \**p* < .05. \*\**p* < .01. \*\*\**p* < .001.

Binomial GLMM results indicate significant main effects of *Block* (χ^2^ = 108.05*, p < .*001), as well as a significant interaction effect of *Session* by *Block* (χ^2^ = 65.10*, p < .*001), whereas no significant main effects of *Session* (χ ^2^ = 0.33*, p* = .85), *chosen side* (χ^2^ = 0.65*, p* = .42), *baiting time* (χ^2^ = 0.02*, p* = .89), and *trial sequence* (χ^2^ = 0.24*, p* = .63) were observed.

To explain the absence of a main effect of *Session*, further post-hoc pairwise comparisons indicate that our subjects demonstrated significantly poorer performance in the test block of the *crossed* session compared to the *baseline* session (difference = 2.38, *SE* = 0.25, *z* = 9.36*, p < .*001) and the *contact* session (difference = 1.79, *SE* = 0.23, *z* = 7.74*, p < .*001; also see Figure 4a). However, there was no significant difference in correct choices between the test block in the *baseline* and *contact* sessions (difference = 0.60, *SE* = 0.26, *z* = 2.30*, p* = .15). Similarly, within these sessions, subjects’ accuracy declined significantly only in the test block of the *crossed* session compared to their performance in the pre-test baseline block (difference = 2.22, *SE* = 0.25, *z* = 9.03*, p < .*001) and the post-test baseline block (difference = 2.05, *SE* = 0.24, *z* = 8.58*, p < .*001). For the other two sessions, no significant variations in choice accuracy between blocks were observed. All p-values were Bonferroni-Holm corrected, and metrics were absolutised.

**Figure 4.**
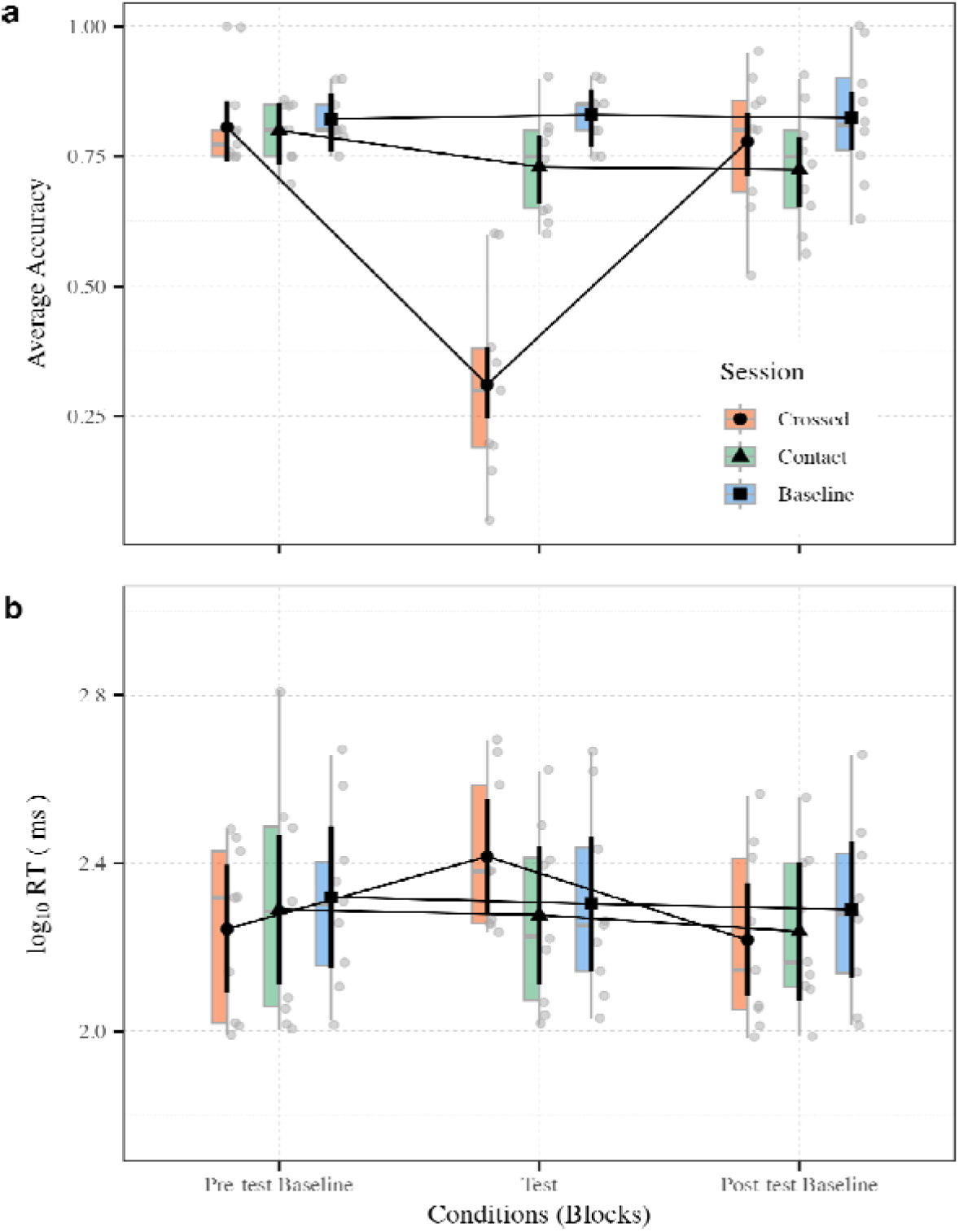
Averaged choice accuracy (aggregated by subject) and Log-transformed RTs plotted as a function of sessions and blocks. *Note:* Coloured points in (a) show the average accuracy of individual subject; in (b) show the RT (measured in *ms*) transformed by taking the base-10 logarithm per trial of individual subjects. Coloured boxplots demonstrate the median and quartiles, black points in the foreground show the mean, error bars show 95% confidence intervals.

### Reaction Time (RT)

GLMM results indicate a significant interaction of *session* by *block*, *F* (4, 1487.84) = 10.28*, p < .*001. Furthermore, we observed significant main effects *block* [*F* (2, 40.03) = 26.26*, p < .*001], as well as *chosen side* [*F* (1, 1501.16) = 4.50*, p* = .034] on log-transformed RT (see Figure 4b). No significant main effects of *session* [*F* (2, 19.69) = 1.36*, p* = .28], *trial sequence* [*F* (1, 1488.69) = 0.85*, p* = .35], and *baiting duration* [*F* (1, 1390.99) = 0.13*, p* = .72], were observed.

Followed-up pairwise comparison results demonstrate a significant longer *log*_10_RT in the test block of *crossed* session than the *contact* session [*difference* = 0.14, *SE* = 0.03, *t*(22.1) = 4.10*, p = .*004], as well as the *baseline* session. However, the latter difference is not statistically significant, *difference* = 0.11, *SE* = 0.05, *t*(13.5) = 2.45*, p* = .229. No significant differences in RTs were observed across other blocks (see Figure 4b).

Furthermore, results showed that the *log*_10_RT only increased significantly in the test block of the *crossed* session compared to the latency in pre-test baseline block, *difference* = 0.17, *SE* = 0.03, *t*(46.4) = 6.09*, p < .*001, and post-test baseline block, *difference* = 0.20, *SE* = 0.03, *t*(36.2) = 6.60*, p < .*001. For the other two sessions, no significant variations in RT between blocks are observed. All p-values were Bonferroni-Holm corrected, and metrics were absolutised.

## Discussion

This study examined the hornbills’ ability and reaction times (RT) in discriminating baited strings across three configurations: *parallel*, *contact*, and *crossed*. These conditions were introduced in a block design, following a sequence of *pre*-test *baseline* block → test block → *post*-test baseline block, forming part of a learning framework. Our data suggests marked differences in string-pulling accuracy and RT across these conditions, shedding light on the cognitive mechanisms employed by large-brain avian species in such problem-solving tasks. These results offer new insights into the goal-directed string-pulling behavior of East Asian hornbills, extending research that has primarily focused on other avian species like passerines and psittacines using the patterned-strings paradigm (Jacobs & Osvath, 2015).

First, our study confirmed that hornbills are capable of reliably performing learned, goal-directed behaviour in the baseline session. Their performance was comparable to that of passerines and psittacines reported in previous studies (e.g., Bagotskaya et al., 2012; Taylor, Medina, et al., 2010; Wakonig et al., 2021). After gaining experience with single- and double-string tasks, all oriental-pied hornbills (*Anthracoceros albirostris*) consistently chose the correct string at rates significantly above chance, without exhibiting a side bias, suggesting that they, like other large-brained avian taxa, can learn and apply associative cues to solve string-pulling problems. Although group-level performance was reliable, two individuals, Harriet, a plain-pouched hornbill (*Rhyticeros subruficollis*), and Pepper, a Papuan hornbill (*Rhyticeros plicatus*), showed a decline in accuracy toward the end of the session, approaching chance levels. This decline is likely due to reduced attention as a result of task duration. In contrast, the Oriental-pied hornbills maintained stable accuracy across blocks. Their consistent performance may reflect a considerable degree of self-control, consistent with findings on their advanced cognitive abilities and adaptability observed in both captive and natural environments (e.g., Garcia-Pelegrin, 2025; Loong et al., 2021; Yao & Garcia-Pelegrin, 2024).

Another challenging condition introduced in our study was the *contact*/*no contact* problem, in which subjects had to distinguish between an intact string and a broken one to obtain the reward. Five out of nine hornbills showed notable flexibility by consistently choosing the intact string and avoiding the broken one at rates significantly above chance. Notably, two of these five successful individuals, Olivia and Oris, developed significant side biases in the subsequent post-test baseline block. This may reflect a temporary depletion of executive control following a cognitively demanding task, as has been documented in humans (Schmeichel, 2007; Schmeichel et al., 2003; Shelton et al., 2011). One subject, Sadie, also developed a significant side bias during the test block, but this appeared idiosyncratic, as she had already shown a bias in the pre-test baseline block.

At the group level, hornbills performed equally well in the *contact* test block compared to the *baseline* test block and both the pre- and post-test *baseline* blocks. Performance in the *contact* condition was also significantly better than in the *crossed* condition. These results are consistent with previous findings in passerines and psittacines, including azure-winged magpies (Wang et al., 2019), Australian magpies (Johnsson et al., 2023), hooded crows (Obozova et al., 2014), Lear’s and hyacinth macaws (Schuck-Paim et al., 2009), and galahs (Krasheninnikova, 2013). Notably, similar success has been observed in northern ground-hornbills (Danel et al., 2019), a species from a different order (*Bucorvidae*) but the same family (*Bucerotiformes*) as our hornbill subjects (*Bucerotidae*), with approximately half of tested individuals solving the *contact* task with relative ease.

The *crossed* condition posed a more complex problem, requiring subjects to recognise and respond to the spatial displacement between the accessible end of the string and the baited reward. None of the hornbills demonstrated an understanding of this spatial relationship. Additionally, 4 out of 9 individuals exhibited a proximity bias, consistently selecting the string end closest to the reward, mirroring their behaviour in the *baseline* condition. Similar proximity-based responses have also been reported in non-human primates (e.g., Albiach-Serrano et al., 2012; Cha & King, 1969), suggesting a widespread reliance on immediate visual-spatial cues across taxa. Another 4 subjects developed a significant side bias, repeatedly choosing strings on one side more frequently than the other, with two individuals maintaining this bias into the subsequent post-test baseline block. This strategy may reflect a secondary, efficiency-driven approach to binary-choice tasks, where consistently selecting one side yields a 50% chance of success without requiring an understanding of the task, and the strategies have been observed in other species when task demands increase or when problems become “unsolvable” (Gagné et al., 2012; Jacobs & Osvath, 2015; Wakonig et al., 2021).

Group-level analysis supported these individual patterns, revealing a significant decline in accuracy during the test block of the *crossed* session, compared to both the pre- and post-test baseline blocks and the test blocks in both the *baseline* and *contact* sessions. The hornbills’ performance aligns with previous findings in passerines and psittacines, where few individuals solve the crossed-string task, and most develop biases towards certain spatial features (e.g., Hofmann et al., 2016; Krasheninnikova, 2013; Schuck-Paim et al., 2009; Taylor, Medina, et al., 2010; Wakonig et al., 2021; Wang et al., 2019). Overall, our results indicate that, under the *crossed* condition, naïve hornbills are more likely to rely on surface-level visual cues than to engage in spatial reasoning about string connectivity (Taylor, Medina, et al., 2010; Taylor et al., 2012). Nevertheless, some individuals exhibited behavioural flexibility by adopting consistent side-based strategies to maximise rewards despite limited task comprehension.

The hornbills’ success in the *contact* problem suggests that they might be attuned to contextual cues during string-pulling, and possibly more adept at recognising physical continuity than tracking string displacement. More importantly, it indicates that they, at least partially, focus on the string configurations rather than simply monitoring the reward’s position, as opposed to the proximity hypothesis in the *crossed* condition (see Jacobs, 2018; Jacobs & Osvath, 2015). Therefore, failures in the *crossed* condition might be more likely to result from difficulty disentangling the physical intertwinement of the strings. This explanation is supported by findings from Krasheninnikova et al. (2013), which showed that parrots solved the crossed-strings problem when the strings were marked with distinct colours, but struggled with unmarked strings.

Analyses of hornbills’ reaction times (RTs) across the three sessions showed that subjects responded significantly more slowly in the *crossed* test block compared to the *contact* test block. Although RTs were also elevated in the *crossed* relative to the *baseline* test block, this difference was not statistically significant. By contrast, RTs in the *contact* and *baseline* test blocks were highly similar. Within-session comparisons further revealed that hornbills took significantly longer to act during the *crossed* test block than during the corresponding pre- and post-test *baseline* blocks. No such RT variation was observed within the *contact* or *baseline* sessions, where reaction times remained consistent across all three blocks. Interestingly, despite some individuals showing reduced accuracy in the post-test *baseline* blocks, group-level RTs remained stable across these blocks in all sessions. This dissociation suggests that performance declines in certain subjects may not reflect general cognitive fatigue or resource depletion, but are instead more likely driven by individual-level factors that do not manifest consistently at the group level.

A traditional perspective on reaction time (RT) is that it reflects the amount of computation involved in perceptual and motor planning before initiating an action (Ratcliff, 1978, 2002). Interestingly, our findings suggest hornbills demonstrated task-complexity mediated response latency, where they spent significantly longer time acting on the strings in a novel and more challenging version that they did poorly (i.e., the *crossed*), compared to a familiar and challenging one (the *contact/no contact*) and an easy one (*baseline*, two-string discrimination). Although we did not anticipate that RTs would directly reflect deliberation in planning under the patterned-strings paradigm, the increased RTs may indicate that hornbills internally processed the *crossed* pattern due to its higher complexity in string alignments. However, their failure to solve the task may be attributed to difficulties in distinguishing physical compartments when no clear visual cues signaled separation, or the prolonged action planning time may not have been sufficient to lead to a correct solution due to less executive control. This interpretation could also align with our findings in the *baseline* and *contact* test blocks, where clear physical separation, either between two strings or within a broken string, led to shorter RTs and comparable accuracy, contrasting with the *crossed* condition.

Beyond the classical interpretation of reaction time (RT) as an index of mental computation, recent research suggests that RT can also be shaped by habituation biases arising from prior exposure to task-relevant spatial or kinematic features during motor learning (Diedrichsen et al., 2010; Verstynen & Sabes, 2011; Wong et al., 2016, 2017). For example, Wong and colleagues (2017) demonstrated that participants’ RTs differed across two otherwise identical tasks depending on their interim training in specific kinematic habits. In their second experiment, RTs remained prolonged even in a simpler, cued version of the task following prior training on a more complex, uncued version, indicating that the slower responses were driven by entrenched motor habits rather than increased cognitive load. Applying this framework to our study, the comparable RTs observed under the baseline (parallel) and contact conditions may reflect the activation of familiar perceptual or motor schemas when the bird encountered continuous string configurations. In contrast, the prolonged RTs observed in the crossed condition likely stem from limited habituation to its novel, non-continuous layout. Although our current data do not directly confirm a habituation bias, particularly as detailed kinematic patterns were not measured, we suggest that future research could incorporate technologies such as infrared motion capture and predictive modelling on movement estimation to more precisely investigate these effects in avian species (e.g., Cronin, 2021; Williams et al., 2021).

Alternatively, the prolonged RTs in the test block of the *crossed* session may result from consecutive failures in a more complex task, in which error-related slowing has been consistently observed in human subjects during some two-alternative forced-choice tasks (e.g., post-error slowing; Danielmeier & Ullsperger, 2011; Debener et al., 2005; Gehring & Fencsik, 2001). Related phenomenon is linked with executive functions in humans and other mammals, such as value updating, conflict monitoring, and inhibitory control, which are critically associated with activities in the neural circuits including the orbitofrontal cortex (OFC), medial frontal cortex (MFC), and basal ganglia (see Ullsperger et al., 2014, for a review). Notably, the neural networks involved, particularly those in the frontal cortex, have been found to show functional analogies to the avian nidopallium caudolaterale (see Rose & Colombo, 2005). As post-error slowing remains under-explored in avian species, we encourage future research utilising further behavioural as well as neurophysiological methods to investigate its potential occurrence.

In addition to our assessments of behaviour across string patterns, we observed a general trend of laterality in RTs, with hornbills responding faster to the string on their left side, regardless of which string was baited. This RT advantage likely stems from brain lateralisation, a phenomenon widely documented across species (for a review, see Güntürkün et al., 2020). Lateralisation is believed to be associated with enhanced cognitive and behavioural flexibility by allowing certain hemispheres to specialise in distinct tasks, improving information processing efficiency (Magat & Brown, 2009; Rogers, 2000; Vallortigara & Rogers, 2005). In birds, this often manifests as side biases in tasks such as retrieval or detection of ecologically relevant stimuli, with one hemisphere typically dominant in sensory and motor control (see Vallortigara et al., 1999). Such lateralisation is also evident in large-brained species like corvids and parrots (e.g., Harris, 1989; Hunt et al., 2001; Rogers, 1980; Wiltschko et al., 2002). Although lateralisation has been well-explored in these birds, research within the order *Bucerotidae* remains limited. Our findings contribute to this body of knowledge, suggesting that lateralisation may be a universal trait in large-brained avian species, despite the small effect size in our data. Future studies using neuroimaging or controlled behavioural experiments could further explore lateralisation in hornbills and its role in their cognitive abilities (Güntürkün et al., 2020).

One limitation of this study is that it was conducted outside a well-controlled laboratory setting. Instead, the subjects were tested in their familiar zoo enclosures, housed adjacent to other wild animals. As a result, random environmental noises were inevitable and may have interfered with the subjects’ perceptual or cognitive processing during testing. However, we managed to mitigate these effects by employing filters for extreme anomalies as well as more liberal statistical methods, accounting for by-subject, trial, and session-level random noise in our models. With that being said, future research could address this limitation by testing in a lab environment or using a digitalised string-pulling task paradigm for maximum control over the experimental procedures (e.g., Wasserman et al., 2013). Furthermore, it is important to note that our results may not generalise to hornbills’ natural temporal tendencies in the *crossed* condition. This limitation stems from the use of a within-subject design within a learning framework, which was intended to maximise data collection given the small sample size (see Farrar et al., 2020). This approach did not differentiate between learning effects and spontaneous knowledge. Therefore, we recommend that future research adopt a between-subjects design and examine either a larger sample of the current species or more accessible avian species with similar behavioural flexibility to investigate the observed effects across avian taxa.

Our study deepens the understanding of goal-directed string-pulling behaviour in East Asian hornbills by exploring their cognitive strategies in problem-solving tasks that range from simple to complex, utilising metrics of accuracy and reaction time. Hornbills exhibited proficiency in simpler configurations, such as the *baseline* condition, and demonstrated success in the more complex *contact* condition. However, they faced significant challenges in understanding the spatial displacement of the *crossed* condition, with some individuals displaying pronounced proximity and side biases. The reaction time data indicate that hornbills experienced prolonged RTs when responding to the *crossed* condition, suggesting they likely processed the *crossed* pattern with greater caution. Nonetheless, it is also possible that a habituation bias to these settings contributed to the observed reaction time disadvantage rather than cognitive computations alone. Additionally, findings of lateralisation in RT data may further emphasise the link between hornbills and advanced cognitive abilities, although the effect size is relatively small. Despite limitations related to environmental control and sample size, our findings pave the way for future investigations into the cognitive mechanisms underlying string-pulling behaviour in hornbills and related avian taxa. By delving deeper into these cognitive processes, we can enrich our understanding of avian intelligence and its broader implications for behavioural ecology.

## Figures

Figures used for data visualisation were produced using R packages “ggplot2” (Wickham, 2016) and “afex” (Singmann et al., 2024).

## Notes

### Competing Interest Statement

The authors have declared no competing interest.

