## Supplementary material A for "Hornbills Adjust Response Speed According to Solvability of Patterned-Strings Problems"

**Table S1**

*Order of sessions tested for each subject*

Subject	1*st* Session (*dd/mm/yy*)	2*nd* Session (*dd/mm/yy*)	3*rd* Session (*dd/mm/yy*)

| Chiku Chika Sadie Sam | Contact (*18/10/23*) Crossed (*01/11/23)* Crossed (*25/10/23)* Crossed (*16/10/23*) | Baseline (*24/10/23*) Contact (*07/11/23*) Contact (*31/10/23*) Baseline (*23/10/23*) | Crossed (*08/11/23*) Baseline (*08/11/23*) Baseline (*01/11/23*) Contact (*14/11/23*) |
| --- | --- | --- | --- |
| Pepper | Contact (*05/12/22*) | Crossed (*12/12/23* A.M.) | Baseline (*12/12/23* P.M.) |
| Oris | Contact (*16/02/24*) | Baseline (*21/02/24*) | Crossed (*23/02/24*) |
| Harriet | Crossed (*07/11/23*) | Baseline (*08/11/23*) | Contact (*15/11/23*) |
| Olivia | Crossed (*27/11/23*) | Contact (*28/11/23*) | Baseline (*11/12/23*) |
| Oscar | Crossed (*10/11/23*) | Contact (*21/11/23*) | Baseline (*22/11/23*) |

*Note.* All data were logged in a paper-based experimental diary. For two sessions conducted in the same test day, the temporal interval in between was at least 4 hours. The order of testing sessions was subjected to food reward allowance per test day.

### **Supplementary material B: Coding Process**

To ensure coding consistency and efficiency, the main experimenter coded all experimental trials (*N* = 1647). For each experimental trial, the we coded the discrete variables: the side of the baited string (L: left vs. R: right), the side of the chosen string (L vs. R), the subject’s choice outcome (correct vs. incorrect); as well as continuous variables: the experimenter’s baiting time (in ms), and the subject’s reaction time (RT; in ms). Baiting time was defined as the duration from centering the subject with the food reward to the moment the experimenter attached the reward to the strings (see Figure 1b). Reaction time (RT) was measured from the moment the subject initiated a peeking action, after observing the experimenter bait the distant container and push the reachable ends toward the enclosure, to the moment the subject first peeked at the chosen string.

**Supplementary Material C: Data Preprocessing**

We processed the coded data files using R package "tidyverse" (Wickham et al., 2019). After processing, the data attributes are:

### **Table S3**

*Attributes retained in the data*

| Attribute code | level/unit | Note |
| --- | --- | --- |
| Subject | *N* = 9 | the name of the subject |
| Session | 3 levels: Crossed, Contact, Baseline | experimental session named after the test block within |
| Condition | 3 levels: Pre-test, Test, Post-test | sequence of blocks within one session |
| BaitedSide | 2 levels: Left, Right | the side of the baited string (in reference to |

the subject)

ChosenSide	2 levels: Left, Right	the side of the string chosen

(in reference to the subject)

| Outcome | 2 levels: correct, incorrect | subject’s decision outcome |
| --- | --- | --- |
| Encode | in *ms* | temporal duration from centering the subject |
| RT | in *ms* | with food reward, until the moment experi- menter attached the reward to the strings  temporal duration from the moment that the |
|  | | subject initiated the peaking action after ob- |
|  |  | serving the experimenter baited and pushed |
|  |  | to the moment that the subjects first peaked |
|  |  | at the chosen string |

We adopted Tukey’s method to detect extreme anomalies in RT per subject by experimental blocks (Tukey, 1977). According to this method, for a univariate distribution, values falling outside the range of [*Q*_1_ *−* 1*.*5 *× IQR, Q*_3_ + 1*.*5 *× IQR*] are considered potential outliers. Here, *Q*_1_ represents the first quartile, *Q*_3_ represents the third quartile, and the *IQR* (inter-quartile range) is defined as *Q*_3_ *− Q*_1_. This method is robust against skewed data, as the RT data we obtained is likely to follow a log-normal distribution (Ulrich & Miller, 1993).
